# Metabolic profiling of drought tolerance: revealing how citrus rootstocks modulate plant metabolism under varying water availability

**DOI:** 10.1101/2022.07.09.499439

**Authors:** Simone F. Silva, Marcela T. Miranda, Camila P. Cunha, Adilson P. Domingues-Jr, Juliana A. Aricetti, Camila Caldana, Eduardo C. Machado, Rafael V. Ribeiro

## Abstract

Water stress is a major environmental factor affecting *Citrus* spp. and Rangpur lime is a drought-tolerant rootstock used to enhance orange yield in rainfed orchards. Here, we combined morpho-physiological analyses with metabolic profiling of roots and leaves of Valencia orange scions grafted onto Rangpur lime, Swingle citrumelo or Sunki mandarin rootstocks under water deficit. Our aim was to present a comprehensive spatio-temporal evaluation of citrus responses to drought and highlight the metabolic adjustments associated with drought tolerance induced by Rangpur lime. Plant responses were evaluated during the initial phase of reduction in water availability, when water deficit was maximum and also after rehydration. Fifty-eight primary metabolites were modulated by water deficit, mainly amino acids, organic acids and sugars. Metabolic changes indicated adjustments related to osmotic, energetic and redox processes under low water availability, which were dependent on rootstock and varied between roots and leaves and along the experimental period. Rangpur lime prioritized root growth in the initial phase of water deficit, which was linked to less sugar accumulation, changes in nucleotide metabolism, downregulation in Shikimic acid pathway and accumulation of arginine. After rehydration, the resume of shoot growth was associated with high accumulation of arginine and asparagine. The better performance of Rangpur lime seems to be associated with its high sensitivity of roots to changes in water availability and possible signaling compounds have been suggested.

## Introduction

Water deficit is the main factor limiting agricultural productivity (Chaves et al., 2003) and uncertain rainfall and warming due to climate change should modify drought intensity, duration and frequency worldwide (IPCC, 2022; Thornton et al., 2014). Understanding how plants respond to drought stress becomes a hot topic worth investigation for promoting a sustainable agriculture in a changing environment (Kang et al., 2009). While citrus orchards are susceptible to seasonal water shortage (Ribeiro et al., 2006; Thornton et al., 2014), the grafting technique has improved the performance of citrus trees under drought in a rootstock-dependent manner (Ribeiro et al., 2014; Martínez-Cuenca et al., 2016).

Among citrus rootstocks, Rangpur lime (*Citrus limonia* Osbeck) is known by its drought tolerance linked to morpho-physiological strategies to maintain shoot water status, leaf gas exchange and plant growth under water deficit (Mourão Filho et al., 2007; Ribeiro and Machado, 2007; Pedroso et al., 2014; Miranda et al., 2018; Bowman and Joubert, 2020; Miranda et al., 2021; Silva et al., 2021). Rangpur lime presents high root hydraulic conductivity and internal hydraulic redistribution (Medina and Machado, 1998; Medina et al., 1998; Miranda et al., 2018; Miranda et al., 2021), besides being able to detect subtle variations in water availability (Santana-Vieira et al., 2016; Miranda et al., 2020; Silva et al., 2021). These traits improve shoot water balance and together with an effective stomatal regulation favor leaf gas exchange under water deficit. In fact, plants need a fine tuning to reduce water loss while maintaining photosynthesis, which modifies growth pattern to balance energy supply and demand under varying water availability (Flexas et al., 2006; Chaves and Oliveira, 2004; Smith and Stitt, 2007). Increased sink demand by photoassimilates and enhanced root growth under water deficit are other two important morpho-physiological responses of Rangpur lime to water deficit (Pedroso et al., 2014; Silva et al., 2021). However, underlying metabolic processes that support the drought tolerance syndrome found in Rangpur lime have not been fully elucidated.

Acclimation to drought involves the reconfiguration of metabolic pathways to establish a new homeostasis, which alters the metabolite profile of plant organs (Shulaev et al., 2008). Under water deficit, the main groups of altered metabolites are those related to energetic balance, compatible osmolytes for maintaining cell turgor or ones to deal with free radicals and then protect cell functions (Goufo et al., 2017; Guo et al., 2018). High levels of proline, activation of polyamine synthesis, activation of sugar synthesis and changes of branched-chain amino acid metabolism (valine/leucine/isoleucine pathway) are common responses in plants under water deficit (Mata et al., 2016). Accumulation of aromatic amino acids intermediate from the shikimate pathway – which have a protective role against ROS and serve as precursors to a wide range of secondary metabolites – has been reported in leaf tissues under low water availability (Bowne et al., 2012; Michaletti et al., 2018; Guo et al., 2020). Decreased levels of organic acids of tricarboxylic acid cycle (TCA cycle) have suggested inhibition of respiratory metabolism under drought (Das et al., 2017; Prinsi et al., 2018; Jia et al., 2020), while changes in nucleotide metabolism have been associated with ATP supply and nitrogen remobilization in drought-stressed plants (Das et al., 2017; Guo et al., 2020). Furthermore, the metabolite profiles differ among plant organs (Zhang et al., 2017, Ullah et al., 2017, Goufo et al., 2017; Kang et al., 2019) and also due to duration and severity of water deficit (Qu et al., 2019; Itam et al., 2020).

Although the metabolic profile has already been used to understand drought tolerance in citrus species (Pérez-Pérez et al., 2007; Zandalinas et al., 2017; Sousa et al., 2022), new insights are only possible if a comprehensive spatio-temporal analysis is performed considering water deficit intensity and duration, plant recovery capacity, plant organs, and modulation by rootstocks in citrus trees. Assuming that the morpho-physiological changes already identified in Rangpur lime are associated with the reconfiguration of metabolic pathways, the identification of key metabolites – and then metabolic pathways – associated with drought tolerance and how such reconfiguration affects scion metabolism become essential to reveal the rootstock modulation of citrus responses to drought and then link micro (cellular level) to macro (plant level) changes under varying water conditions. Herein, we combined morpho-physiological responses and the primary metabolite profiling of roots and leaves to reveal how Valencia orange scions grafted on three rootstocks respond to varying water availability. In fact, our results confirmed the better performance of plants grafted on Rangpur lime under water deficit, but we now describe and compare the metabolic pattern among three citrus rootstocks. Then, the metabolic alterations found only in Rangpur lime were highlighted as well as the metabolic markers of drought tolerance, which may further assist on the development of drought-tolerant citrus species.

## Material and methods

### Plant material and growth condition

Ten-month old Valencia orange [*Citrus sinensis* (L.) Osbeck] scions grafted on Swingle citrumelo [SC, *Citrus paradisi* Macf x *Poncirus trifoliata* (L.) Raf.], Rangpur lime (RL, *Citrus limonia* Osbeck) and Sunki mandarin (SM, *Citrus sunki* Hort. ex Tanaka) rootstocks were grown in plastic bags (4.5 L) containing organic substrate composed of pine bark (Tropstrato V8 Citrus, Vida Verde, Mogi Mirim SP, Brazil), under greenhouse conditions. We pruned the main trunk (∼ 10 cm cut off from the apex) to stimulate sprouting of new branches and standardize plant height and leaf area before moving plants to a growth chamber (PGR15, Conviron, Winnipeg, Canada) with 30/20 °C (day/night), 12-hour photoperiod (7h00-19h00), photosynthetically active radiation (Q) of 800 μmol m^-2^ s^-1^, and air vapor pressure deficit (VPD) lower than 2 kPa. After an eight-day acclimation period, plants had three young sprouts (∼ 0.5 cm).

### Water regimes

Five well-watered plants of each rootstock were initially evaluated (time-point 0, *n* = 5) and other 30 plants were grouped into two sets (*n* = 15), one maintained well-watered (control) and the other subjected to water deficit (WD). Control plants were grown with substrate moisture at ∼80% of the maximum water storage capacity (MWC). The period under WD treatment was classified in three phases: (i) phase I, slow and initial reduction in water availability by water withholding until the substrate reaches ∼25% MWC [18 days after the first evaluation (DAE)]; (ii) phase II, water deficit by maintaining the substrate at ∼25% MWC for 15 days (from 18 to 33 DAE); and (iii) phase III, substrate rehydration by increasing and maintaining MWC at ∼80% (from 33 to 39 DAE) (**Supplementary Fig. S1**). Substrate moisture was monitored by weighing each bag on a daily basis with an electronic scale, and water was supplied when needed to keep the moisture level within the desired levels (Pedroso et al., 2014). Plant sampling was conducted at the end of each phase (18, 33 and 39 DAE).

### Leaf gas exchange and leaf water potential

Leaf CO_2_ assimilation (*A*), stomatal conductance (*g*_s_) and transpiration (*E*) were measured between 8h00 and 11h00 using an infrared gas analyzer (LI-6400F, LI-COR, Lincoln NE, USA), under air CO_2_ concentration (∼ 400 µmol mol^-1^) and photosynthetically active radiation (PAR) of 800 µmol m^-2^ s^-1^ in previously formed (phase I) or new (phases II and III) fully expanded leaves. This change along the experimental period aimed to avoid self-shading interference on leaf gas exchange. Leaf water potential (Ψ) was evaluated between 11h00 and 12h00 using a pressure chamber (model 3005, Soil Moisture Equipment Corp., Santa Barbara CA, USA).

### Relative growth rate and biometry

Main trunk, mature leaves, new shoots (referred here as young sprouts, including young leaves and branches developed during the experimental period) and roots were dried separately in an oven with forced air circulation at 60°C, until constant weight. The relative growth rate (RGR) of new shoots and roots were calculated following Beadle (1993), using dry matter sampled along the experimental period (**Supplementary Fig. S3)**. Number of leaves was counted, main trunk and branch length was measured with a ruler, main trunk diameter was measured with a digital caliper and the total leaf area was assessed with a digital planimeter (LI-3000, LI-COR, Lincoln NE, USA).

### Starch content

Leaves (previously formed or new ones as described for leaf gas exchange) and roots were harvested at phases I to III (18, 33 and 39 DAE) from control and WD plants. Plant material was ground in liquid nitrogen and then lyophilized for measuring starch content in 10-mg samples, according to the enzymatic method proposed by Amaral et al. (2007). Sample absorbance at 490 nm was measured with a spectrophotometer (Genesys 10S UV-Vis, Thermo Fisher Scientific, Waltham MA, USA) and starch concentration determined using a standard curve.

### Metabolic profiling

Polar metabolites were extracted from 15 mg (leaves) or 30 mg (roots) of fresh plant material ground in liquid nitrogen. Pooled samples for each tissue were used to certify the equipment stability throughout the analysis as quality control. Metabolite extraction followed the protocol described by Salem et al. (2016), using methyl tertiary-butyl ether (MTBE):methanol:water (3:1:1, v/v/v). 150 μL-aliquot of the polar phase was vacuum dried at 4°C and stored at -80°C. Derivatization reaction was carried out in two steps as described by Lisec et al. (2006). The samples were reacted for 90 min at 30 °C with 10 μL of 40 mg mL^-1^ methoxyamine hydrochloride in pyridine, followed by 30 min at 37 °C with 90 μL of N-Methyl-N-(trimethylsilyl)trifluoroacetamide (MSTFA). In this step, 13 Fatty Acid Methyl Esters (FAMEs, C8-C30) were added to each sample, including the technical control samples (blank).

An aliquot of 1 μL of each sample was analyzed by high-throughput gas chromatography with time-of-flight mass spectrometer (GC-TOF-MS Pegasus HT, Leco, Saint Joseph MI, USA) equipped with Combi-PAL autosampler and Agilent 7890A gas chromatograph (Agilent Technologies, Waldbronn, Germany), in both split (1:30) and split less modes. The gas chromatographic analysis was performed on a 30 m x 0.32 mm x 0.25 μm DB-35MS (Agilent J&W, Santa Clara CA, USA). The injection temperature was 230°C, with a flow of 2 mL min^-1^ helium, starting after 180 s from the beginning of data acquisition. The initial temperature of the column was 85°C, kept for 2 min, and increased at 15°C min^-1^ until reaching 360°C. The column eluate was introduced into an electron ionization source (electron impact) of GC-TOF-MS. The source temperature was 250°C, the electron beam current was 70 eV, and 20 s^-1^ spectra in the range of m/z 85-500.

Chromatograms were exported to the Leco ChromaTOF-GC software (v. 4.51.6.0, Saint Joseph MI, USA) in which we performed baseline correction and conversion of data to the netCDF file. Deconvolution, determination of retention index, correction of retention time, peak alignment, and identification of metabolites were performed using TargetSearch package (Cuadros-Inostroza et al., 2016) in R-software (www.r-project.org). FAMEs were used as indexes for peak identification and metabolome profiling performed using an *in-house* library. Manual curation (using the ChromaTOF-GC software visualization tool) confirmed the identity of metabolites with low peak apex intensities detected by the TargetSearch. Then, metabolites were quantified by peak intensity of a selective mass and the intensity of each metabolite was normalized by the sample dry weight, the total ion chromatogram (TIC) of each sample, and by the overall replacement of outliers. Finally, values obtained for each metabolite were normalized by dividing each raw value by the median within each metabolite. The final value corresponds to the relative intensity of each identified metabolite (**Supplementary Tables S1 to S4**).

### Data analyses

Leaf gas exchange, leaf water potential, plant biometry and starch content data were analyzed using Bayesian statistics and the JASP software (https://jasp-stats.org), with water regimes and rootstocks as sources of variation. When significant effects of the water regimes or rootstocks were detected, mean values (*n* = 5 or 4) were compared using Bayes Factor (BF_10_). Bayes Factors as evidence for the alternative hypothesis (H_1_) followed Miranda et al. (2021), with positive support to H_1_ when 3<BF_10_<20, and strong support when BF_10_>20.

Metabolites significantly (Tukey test, p<0.05) affected by treatments and with fold-change (FC) lower than -0.5 or higher than 0.5 (WD/control) were shown in heatmaps. Heatmaps show log^2^ FC of normalized mean intensities (*n* = 4) and were generated using Complex heatmap (Gu, 2016) and Circlize (Gu, 2014) packages from R-software (v.3.5.3). Principal Component Analysis (PCA) was performed using the MetaboAnalyst 4.0 (Chong et al., 2019).

## Results

### Early physiological responses induced by Rangpur lime rootstock under drought

Low water availability caused decreases in leaf water potential (Ψ) on all three rootstocks (**Fig. 1a-f**). Regarding leaf gas exchange, Rangpur lime was the first rootstock to respond to WD, with early reduction in leaf CO_2_ assimilation and stomatal conductance in comparison with Swingle citrumelo and Sunki mandarin (**Fig. 1g-l**). Upon rehydration (phase III), control and WD plants resumed initial leaf water status and gas exchange, with Swingle citrumelo inducing faster recovery of photosynthesis than the other rootstocks (**Fig. 1g-i**).

**Fig. 1.**
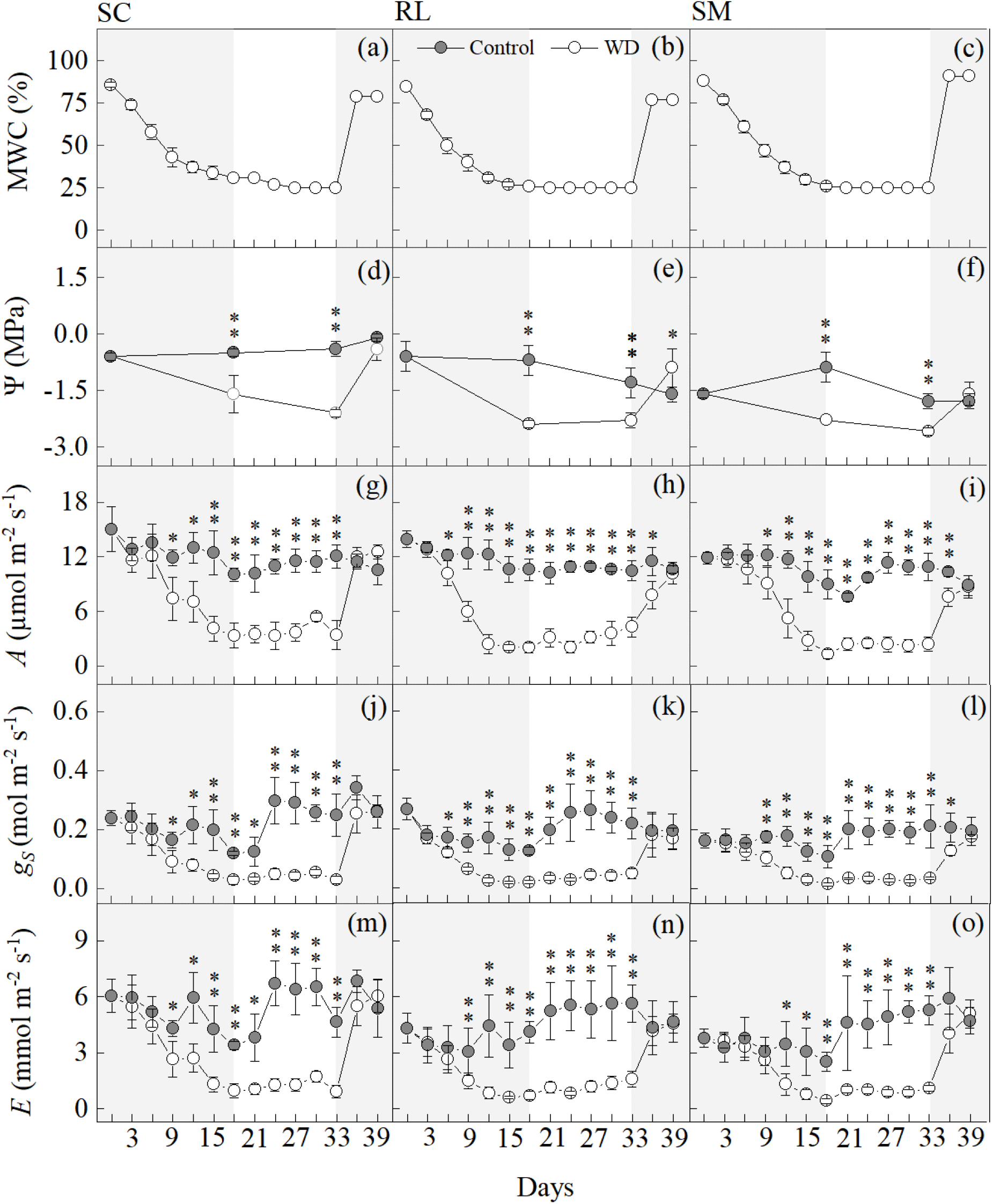
Substrate moisture (a-c) and leaf water potential (d-f), leaf CO_2_ assimilation (g-i), stomatal conductance (j-l) and transpiration (m-o) of Valencia orange scions grafted on Swingle citrumelo (a,d,g,j,m), Rangpur lime (b,e,h,k,n) or Sunki mandarin (c,f,i,l,o) under varying water availability. Control refers to well-watered plants while WD to water-stressed plants. MWC is the maximum water storage capacity of the substrate. Symbols represent mean values (*n* = 5) ± SD. * and ** indicate statistical differences between water treatments at 3<BF_10_<20 and BF_10_>20, respectively. The first shaded area represents phase I of water deficit while the second shaded area represents phase III (rehydration). Between them, phase II is shown.

### Plant growth under water deficit is dependent on citrus rootstock

Changes in number of leaves, leaf area and shoot length during the experimental period revealed that plants grafted on Rangpur lime had reduced growth during the phase I of water deficit (**Fig. 2b,e,h**). Swingle citrumelo responded to low water availability only at phase II (**Fig. 2a,d,g**). Sunki mandarin showed an intermediate response, with reduced number of leaves and shoot length at phase I (**Fig. 2c,i**). After rehydration (phase III), control and WD plants grafted on Swingle citrumelo showed similar increment in number of leaves and shoot length. Plants grafted on Sunki mandarin showed similar increment only in shoot length, regardless the previous exposure to water deficit. Even showing recovery in phase III, plants grafted on Rangpur lime still had reduced number of leaves, leaf area and shoot length under WD as compared with well-watered plants (**Fig. 2b,e,h**). The dry matter of main trunk and previously formed leaves presented non-significant variations due to low water availability (**Supplementary Fig. S2**).

**Fig. 2.**
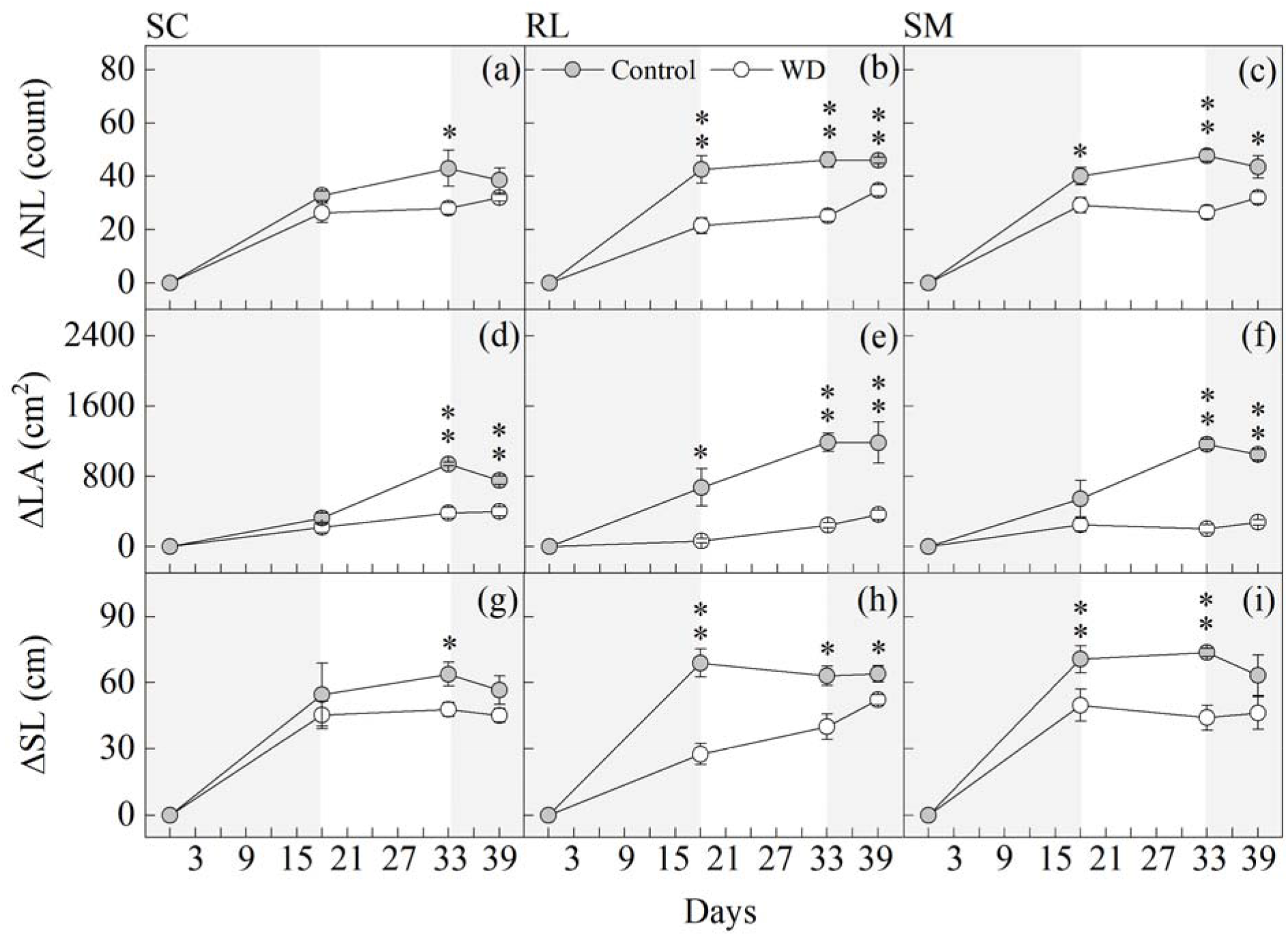
Variation of the number of leaves (ΔNL, a-c), leaf area (ΔLA, d-f) and shoot length (ΔSL, g-i) of Valencia orange scions grafted on Swingle citrumelo (a,d,g,j,m), Rangpur lime (b,e,h,k,n) or Sunki mandarin (c,f,i,l,o) in each phase of water deficit cycle. Control refers to well-watered plants while WD to water-stressed plants. Symbols represent mean values (*n* = 4) ± SD. * and ** indicate statistical differences between water treatments at 3<BF_10_<20 and BF_10_>20, respectively. The first shaded area represents phase I of water deficit while the second shaded area represents phase III (rehydration). Between them, phase II is shown.

Rangpur lime changed plant growth and shoot architecture earlier than the other rootstocks due to water deficit, with a significant reduction in shoot growth and increased root growth at phase I (**Fig. 3a,d**). This pattern of Rangpur lime was changed at phase II, when shoot and root growth under water deficit was similar to one found in control plants (**Fig. 3b,e**). In phase III, there was a significant increase in shoot growth in plants grafted on Rangpur lime and previously exposed to water deficit as compared with well-watered plants (**Fig. 3c**). Sunki mandarin and Swingle citrumelo showed reductions in shoot growth due to water deficit at phase II (**Fig. 3b**), with non-significant differences in plant growth due to previous water deficit being noticed at phase III for both rootstocks (**Fig. 3c,f**).

**Fig. 3.**
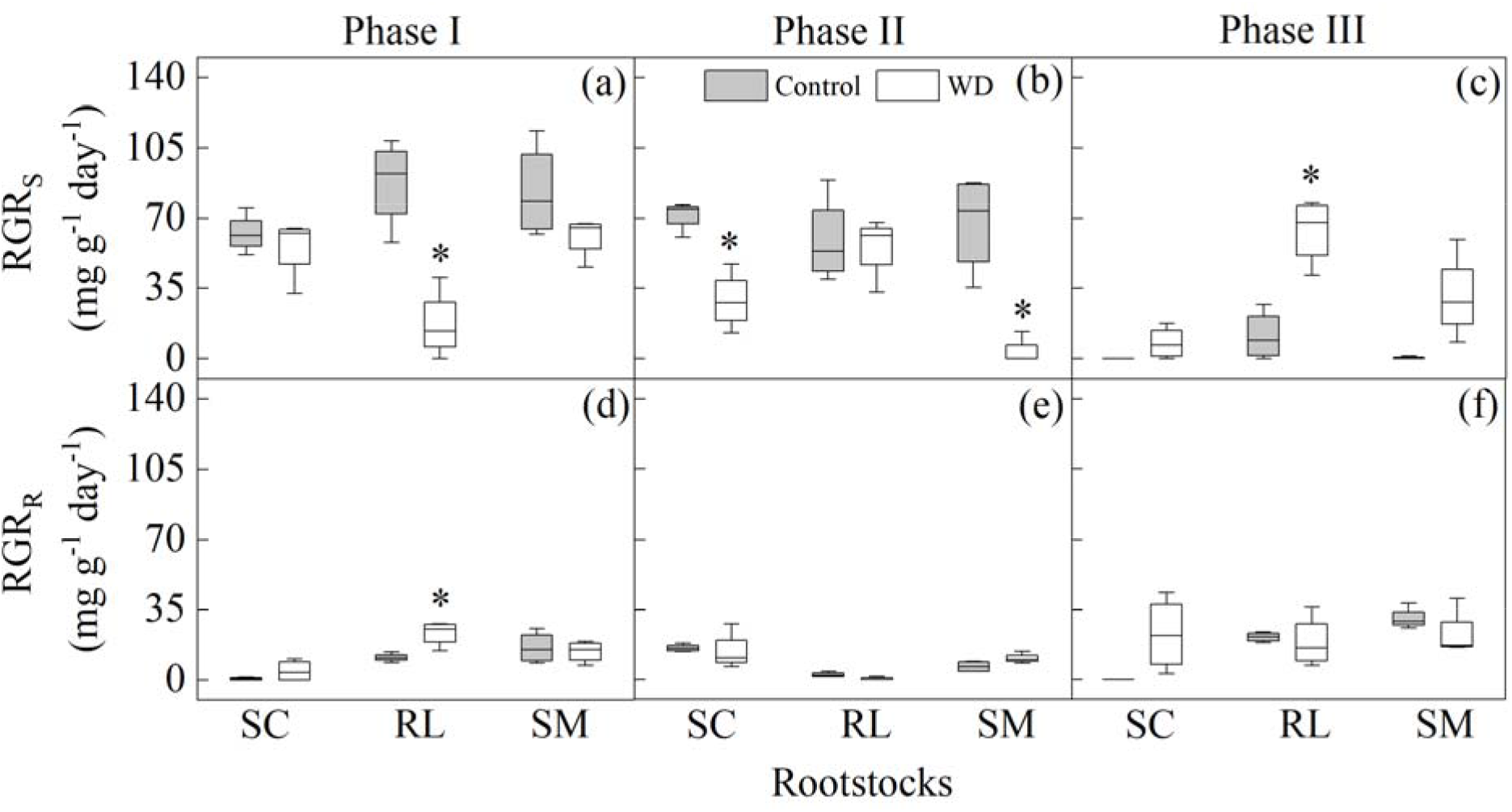
Relative growth rate of shoots (RGRs, a-c) and roots (RGRr, d-f) of Valencia orange scions grafted on Swingle citrumelo (SC), Rangpur lime (RL) or Sunki mandarin (SM) in each phase of water deficit cycle: phase I (a,d); phase II (b,e); and phase III (c,f). Control refers to well-watered plants while WD to water-stressed plants. RGRs is based on dry biomass of new shoots (branches + young leaves). Boxplots consider the median, the 25th and 75th percentiles, the error bars indicate range within 1.5 IQR (*n* = 4). * and ** indicate statistical differences between water treatments at 3<BF_10_<20 and BF_10_>20, respectively.

### Metabolites identified in citrus leaves and rootstocks under water deficit

Metabolic profiling by GC-TOF-MS detected automatically 58 out of 113 polar metabolites in the *in-house* library, considering both leaves and roots (**Supplementary Table S5**). In total, we identified 54 and 45 metabolites modulated by water deficit in roots and leaves, respectively, including mainly amino acids, organic acids, sugars and its derivatives (∼ 90%).

The first two components of PCA explained most of the total variance in roots (between 59% and 80%) and leaves (between 63% and 90%), with PC1 separating samples according to the water regime (**Figs. 4 and 5**). In roots, plants grafted on Rangpur lime were discriminated by arginine and citrulline in all phases (**Fig. 4**). Rootstocks under water deficit were clearly discriminated by sugars at phase I, including erythrose, maltose, trehalose, fructose and glucose (**Fig. 4a**). Differences between water regimes were less evident in roots at phase II (**Fig. 4b**). At phase III, the metabolic divergence between control and water-stressed plants grafted on Rangpur lime was even more evident than at the previous phases, with the contrary being found for the other rootstocks (**Fig. 4c**). In leaves, arginine was responsible for a major distinction between control and water-stressed plants grafted on Rangpur lime at phase I (**Fig. 5a**). Higher discrimination between control and water deficit conditions was found for leaves of plants grafted on Sunki mandarin, which was associated with fructose, fumarate and maleate at phase I and proline and raffinose at phase II (**Fig. 5a,b**). Again, the largest metabolic divergence between control and water deficit samples was found for leaves of plants grafted on Rangpur lime at the recovery period (phase III), with this rootstock being discriminated from the others due to asparagine (**Fig. 5c**).

**Fig. 4.**
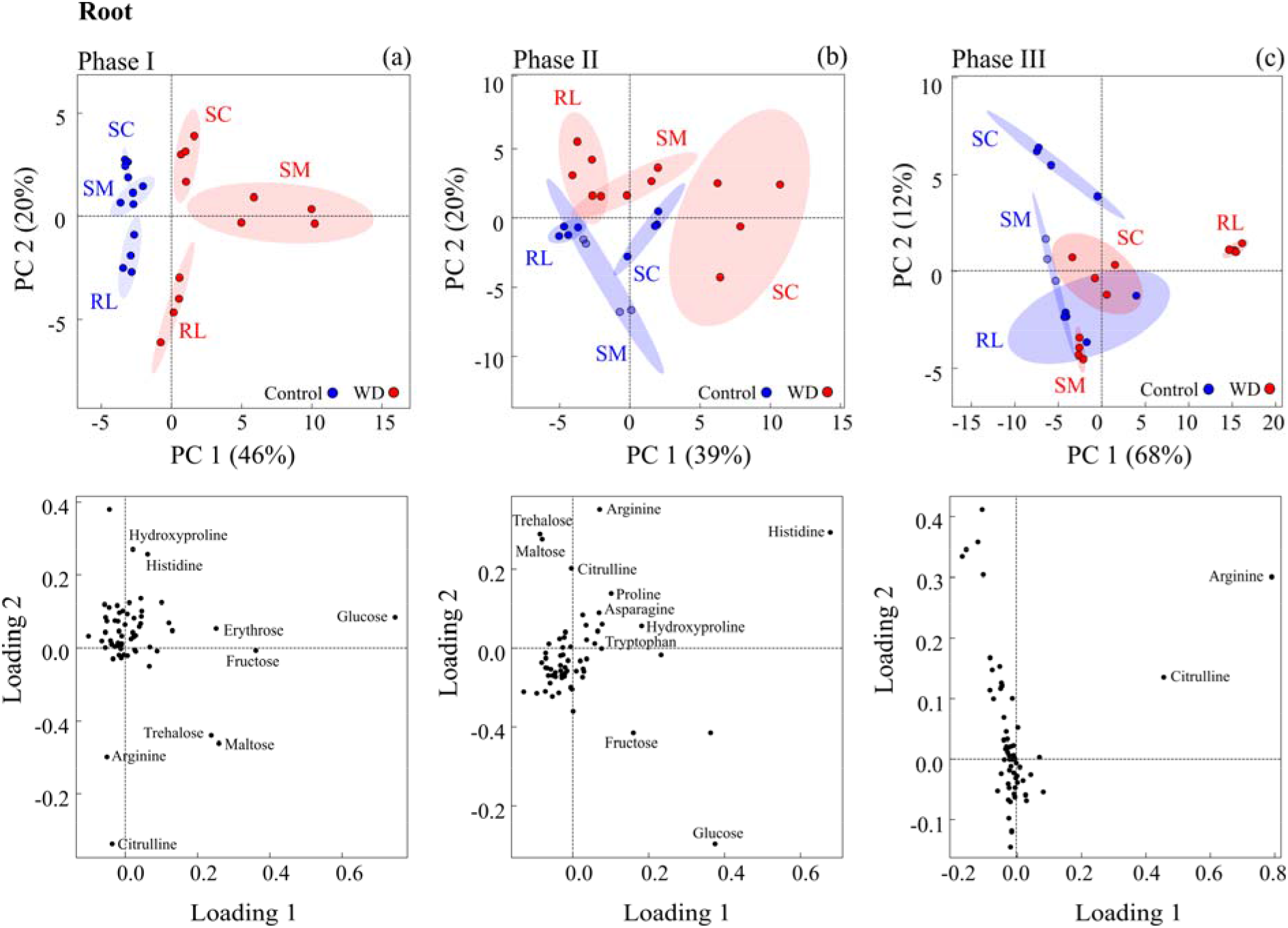
Principal component analysis (score plot and loading plot) of the relative concentration of metabolites identified in roots of Swingle citrumelo (SC), Rangpur lime (RL) or Sunki mandarin (SM) under water deficit (red) and well-watered (blue) conditions. Samples were collected at phases I (a), II (b), and III (c) of water deficit. 95% confidence ellipses are shown.

**Fig. 5.**
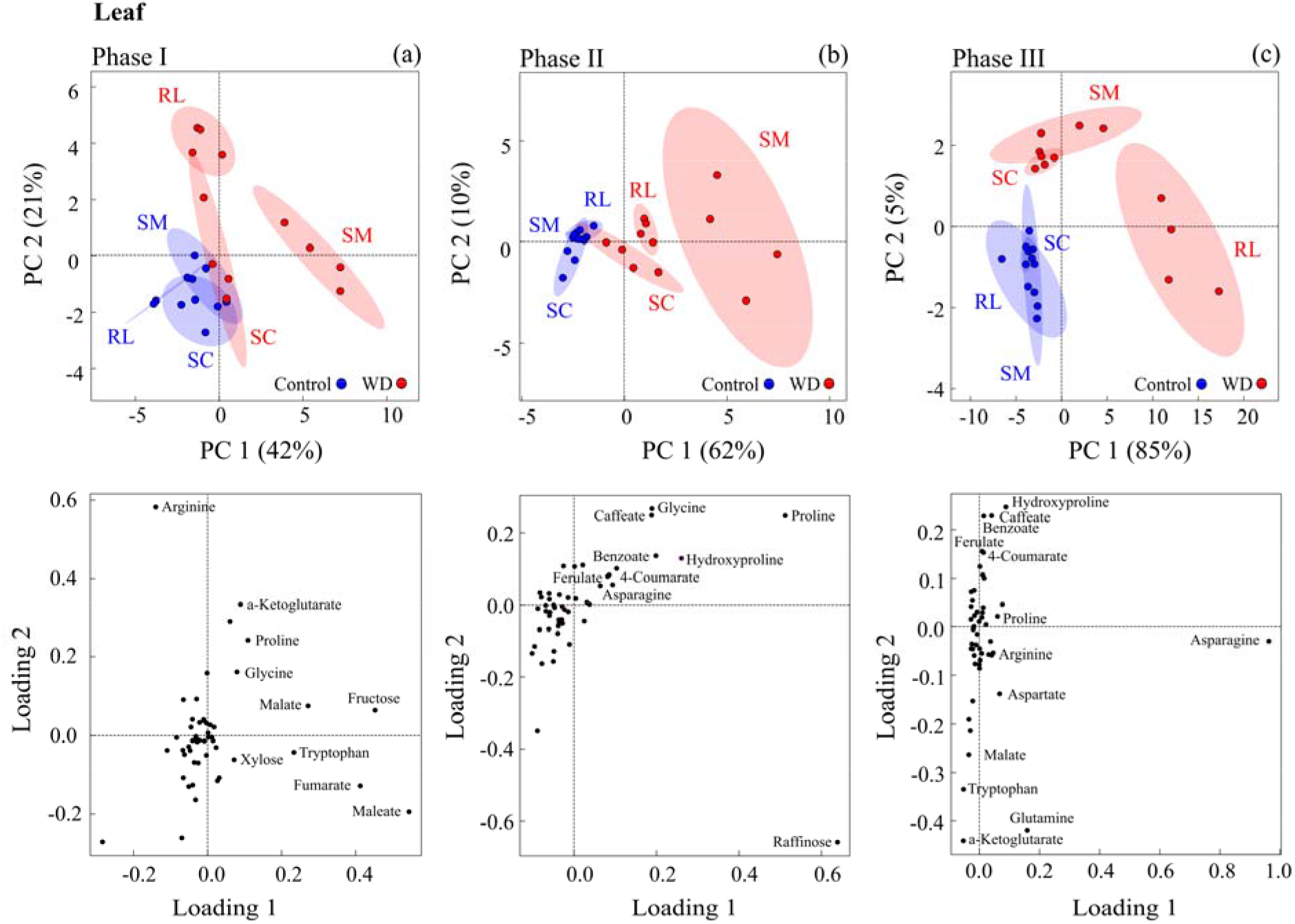
Principal component analysis (score plot and loading plot) of the relative concentration of metabolites identified in leaves of Valencia orange scions grafted on Swingle citrumelo (SC), Rangpur lime (RL) or Sunki mandarin (SM) under water deficit (red) and well-watered (blue) conditions. Samples were collected at phases I (a), II (b), and III (c) of water deficit. 95% confidence ellipses are shown.

### Metabolic profiling altered by citrus rootstocks under water deficit

In roots, there is a clear difference between phases I-II and III, as highlighted by cluster analysis (cluster columns) that identified nine groups of metabolites (**Fig. 6**). The cluster A shows amino acids arginine, citrulline, asparagine, histidine, hydroxyproline, glutamine and proline, which were upregulated at all phases of water deficit, and the sugars maltose, trehalose upregulated during phases I and II. In the cluster B, the sugars glucose and fructose were accumulated during phases I and II of water deficit but downregulated upon rehydration (phase III). The clusters E and F show the downregulation of the organic acids that are intermediates of the glycolysis and TCA cycle (i.e. pyruvate, lactate, α-ketoglutarate and fumarate) in all phases of water deficit. In the cluster G, the levels of amino acids phenylalanine, valine, leucine and isoleucine were reduced during phases I and II but upregulated upon rehydration. Rangpur lime differed from the other rootstocks mainly at phase I, showing more strongly reductions in levels of phenylpropanoids (cluster H).

**Fig. 6.**
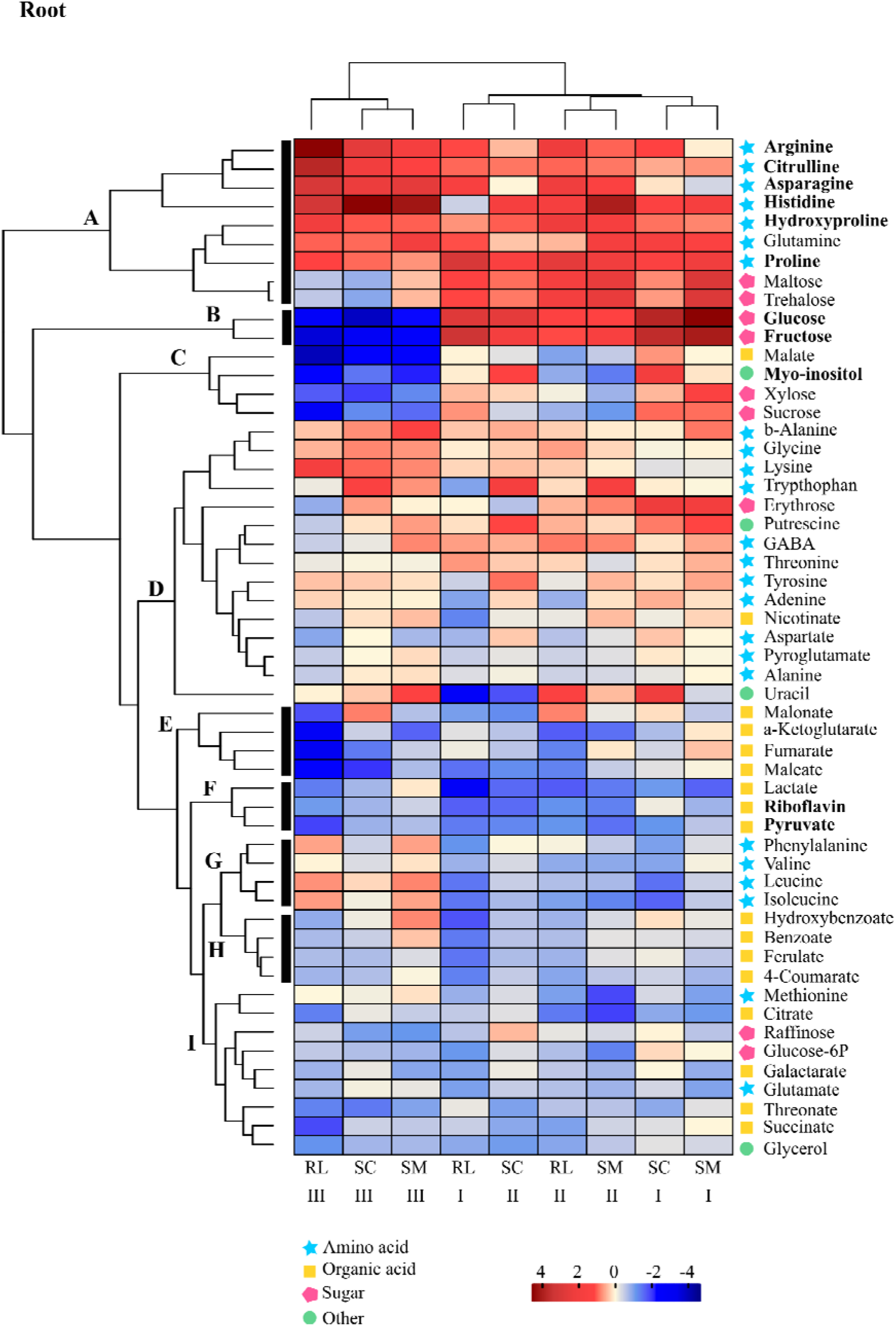
Metabolites identified by GC-TOF-MS in roots of Swingle citrumelo (SC), Rangpur lime (RL) or Sunki mandarin (SM) under water deficit. Samples were collected at phases I, II, and III of water deficit. Heatmap was built using log_2_ FC (water deficit/control) of metabolites. In scale, blue indicates lower and red higher relative concentration. Metabolites identified as drought markers are marked in bold. Capital letters indicate clusters cand black bars highlight the main groups of metabolites.

Regarding leaves, the cluster analysis defined five groups of metabolites (**Fig. 7**). The cluster A contains the amino acids hydroyproline, proline and glycine upregulated at all phases, and asparagine upregulated during the phases II and III. Also in the cluster A, we found phenylpropanoids (i.e. caffeate, benzoate, ferulate and 4-coumarate), which were downregulated at phase I and upregulated at phases II-III, and raffinose upregulated at phase II and downregulated upon rehydration (phase III). The cluster B showed metabolites accumulated at phase I and downregulated later, including TCA organic acids (α-ketoglutarate and malate) and amino acids (tryptophan). The cluster C grouped metabolites accumulated at phase I (e.g. maleate, fructose, fumarate, glutamine and arginine). In leaves, the clear differential response between phases I and II-III demonstrates that the metabolic changes differed between previously formed (phase I) and new (phases II and III) leaves.

**Fig. 7.**
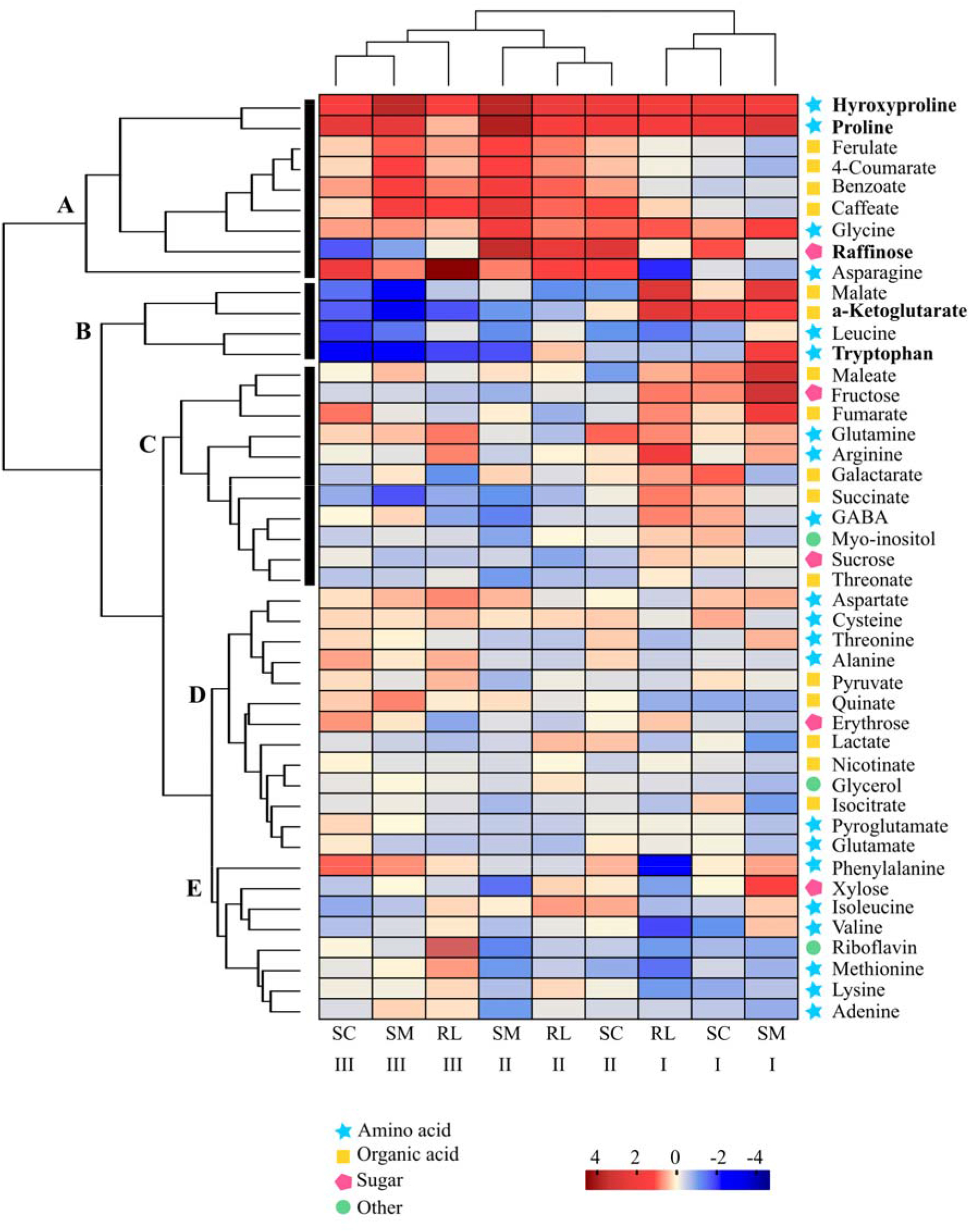
Metabolites identified by GC-TOF-MS in leaves of Valencia orange scions grafted on Swingle citrumelo (SC), Rangpur lime (RL) or Sunki mandarin (SM) under water deficit. Samples were collected at phases I, II, and III of water deficit. Heatmap was built using log_2_ FC (water deficit/control) of metabolites. In scale, blue indicates lower and red higher relative concentration. Metabolites identified as drought markers are marked in bold. Capital letters indicate clusters cand black bars highlight the main groups of metabolites.

Regardless the rootstock, leaf starch concentration was reduced by water deficit at phases II and III (**Supplementary Fig. S4a-c**). However, changes in root starch concentration were dependent on rootstock at phase I, with Sunki mandarin showing reduced starch concentration while Rangpur lime and Swingle citrumelo presenting non-significant changes (**Supplementary Fig. S4d-f)**.

### Metabolite markers for water deficit

Metabolite markers related to drought and rehydration were considered those that show a similar trend among all rootstocks, as well as a high modulation level, given by log_2_ FC water deficit/control higher than 1 or lower than -1 (**Supplementary Tables S6 and S7**). Proline was induced at phases I and II in both roots and leaves, while the accumulation of hydroxyproline was mainly associated with leaves at all phases. According to the phase of water deficit, other potential metabolic markers for roots were: glucose and fructose, both upregulated at phase I; pyruvate and riboflavin (both downregulated) at phase II; and glucose, fructose and myo-inositol (all downregulated) and arginine, citrulline, asparagine and histidine (all upregulated) at phase III. In leaves, the other potential markers were: α-ketoglutarate, upregulated at phase I and downregulated at phase III; raffinose (upregulated) at phase II and tryptophan (downregulated) at phase III. As specific metabolic markers for Rangpur lime, our data revealed arginine and citrulline (upregulated) in roots and arginine (upregulated at phase I) and asparagine (upregulated at phase III) in leaves.

## Discussion

Morpho-physiological and metabolic adjustments of Valencia orange scions grafted onto three rootstock species were evaluated not only under progressive and controlled drought but also after rehydration. Plants grafted on Rangpur lime showed earlier stomatal closure due to the initial decline of substrate moisture as compared to ones on Swingle citrumelo and Sunki mandarin (**Fig. 1j,k,l**). Interestingly, the initial decline in substrate moisture also stimulated Rangpur lime root growth, a morphological strategy to improve water uptake that is supported by enhanced photoassimilate supply to roots under drought (Silva et al., 2021). Overall, Rangpur lime roots were more sensitive and responded rapidly to water deficit, anticipating morphological and physiological reactions before a more drastic reduction in water availability, as described in previous studies (Santana-Vieira et al., 2016; Miranda et al., 2020; Silva et al., 2021). Despite a significant inhibition of growth at phase I, shoot growth resumed at phase II and was maintained at high rates during rehydration (**Fig. 3**), revealing Rangpur lime capacities for acclimation and recovery when facing water deficit.

### Metabolic profiling of citrus plants during and after water deficit

Based on the metabolic profiling, our data indicated adjustments related to osmotic, energetic and redox processes under low water availability, and the importance of amino acid metabolism during rehydration, which were all dependent on rootstock and varied between roots and leaves and along the experimental period (**Figs. 6 and 7**).

Sugar levels increased significantly in roots (mainly glucose, fructose, phase I) and leaves (mainly raffinose, phase II) of all rootstocks under water deficit (**Figs. 6 and 7**). Trehalose – an osmoprotective disaccharide with the ability to stabilize dehydrated membranes (Golovina et al., 2009; Luo et al., 2010) – was also accumulated in roots of all rootstocks, reinforcing previous observation (Santana-Vieira et al., 2016). Unlike previous studies reporting higher raffinose in roots and leaves (Santana-Vieira et al., 2016; Sousa et al., 2022), raffinose accumulation was found only in leaves, mainly young leaves under water deficit (**Figs. 5b and 7**). In our study, starch content was low in these leaves during water deficit (**Supplementary Fig. S4a-c**), which may reinforce the role of raffinose as carbon source during water stress, in addition to its antioxidant action (ElSayed et al., 2013). After rehydration, sugar accumulation in roots and leaves was reversed and decreased sugar levels suggest rapid consumption, probably for generating energy and rebalancing metabolic processes (Peters et al., 2007).

Decreases in leaf CO_2_ assimilation and accumulation of sugars in plants under water deficit (**Figs. 1g,h,i**, **6** and **7**) suggest a downregulation of the respiratory pathway, which is corroborated by reduction in TCA cycle intermediates levels either in roots or leaves – a phenomenon already reported (Prinsi et al., 2018; You et al., 2019; Jia et al., 2020). However, such responses seem to be dependent on organ development stage. Here, the levels of TCA intermediates in roots and new leaves of citrus trees were reduced during the water deficit and rehydration; however, such intermediates were accumulated in previously formed leaves at phase I of water deficit, regardless the rootstock (**Figs. 6 and 7**). Although less common, such accumulation of organic acids was also found in wheat plants under drought (Kang et al., 2019). Changes in the TCA cycle intermediates citrate, malate, α-ketoglutarate, succinate, and fumarate indicate an adjustment in the respiratory metabolism in response to water deficit. Under low carbohydrate availability, branched-chain amino acids (BCAAs), valine, leucine, isoleucine, lysine and phenylalanine can be degraded and used as alternative carbon sources in TCA cycle (Taylor et al., 2004; Araújo et al., 2011). Considering the reduction of BCAAs during water stress, followed by their accumulation after rehydration mainly in roots, we would argue that all rootstocks consumed alternative substrates during water stress (**Figs. 6 and 7**). Alternatively, changes in TCA cycle intermediates may be associated with nitrogen and amino acid biosynthetic pathways.

Phenylpropanoid levels increased significantly in young leaves (phase II), regardless of the rootstock (**Fig. 7**). Phenylpropanoid accumulation has already been reported in leaves of citrus plants under drought (Zandalinas et al., 2017). Herein, we emphasize that such accumulation occurs exclusively in young leaves (developing leaves), which are also more exposed to light. These metabolites are precursors of lignin or can be directly linked to the cell wall and changes in their levels are associated with antioxidant activity and cell wall stiffness (Boba et al., 2017). Higher phenolic content improves cell wall mechanical properties, minimizing water loss while preventing cell elongation (Wang et al., 2016). In fact, accumulation of phenolic compounds in young leaves under water deficit may be associated with leaf expansion inhibition, reduced cuticular transpiration and photoprotection (Hura et al., 2012; Wang et al., 2016).

Proline levels increased significantly in roots and leaves during water deficit, regardless of the rootstock. While proline has a role on osmotic adjustment and redox buffering, hydroxyproline can stabilize the cell wall and provide mechanical resistance against drought (Suguiyama et al., 2014). After rehydration, proline and hydroxyproline accumulation in roots and leaves was reduced.

Higher levels of arginine, citrulline, asparagine and histidine were found in the roots and only asparagine in the young leaves of all rootstocks under water deficit. And, after rehydration the levels of these amino acids were strongly increased (**Figs. 6 and 7**). Besides their osmoprotective function, higher levels of amino acids have been associated with autophagy, proteolysis and restriction of protein synthesis, which favors the supply of precursors for the synthesis of new proteins and secondary metabolites (Hildebrandt, 2018; Prinsi et al., 2018; Batista-Silva et al., 2019). Amino acid accumulation during rehydration may occur due to protein degradation needed for plant recovery after a stressful event (Lyon et al., 2016). In addition, amino acids would act as organic nitrogen reserves, highlighting the importance of nitrogen metabolism during rehydration, a topic to be explored in further studies.

### Underlying processes leading to drought tolerance of Rangpur lime

Unlike the other rootstocks, Rangpur lime showed less sugar accumulation, differentiated changes in nucleotide metabolism (uracil and adenine), downregulation in Shikimic acid pathway and accumulating compounds with high N/C ratio (arginine and asparagine) in roots and leaves, mainly after rehydration (**Fig. 8**).

**Fig. 8.**
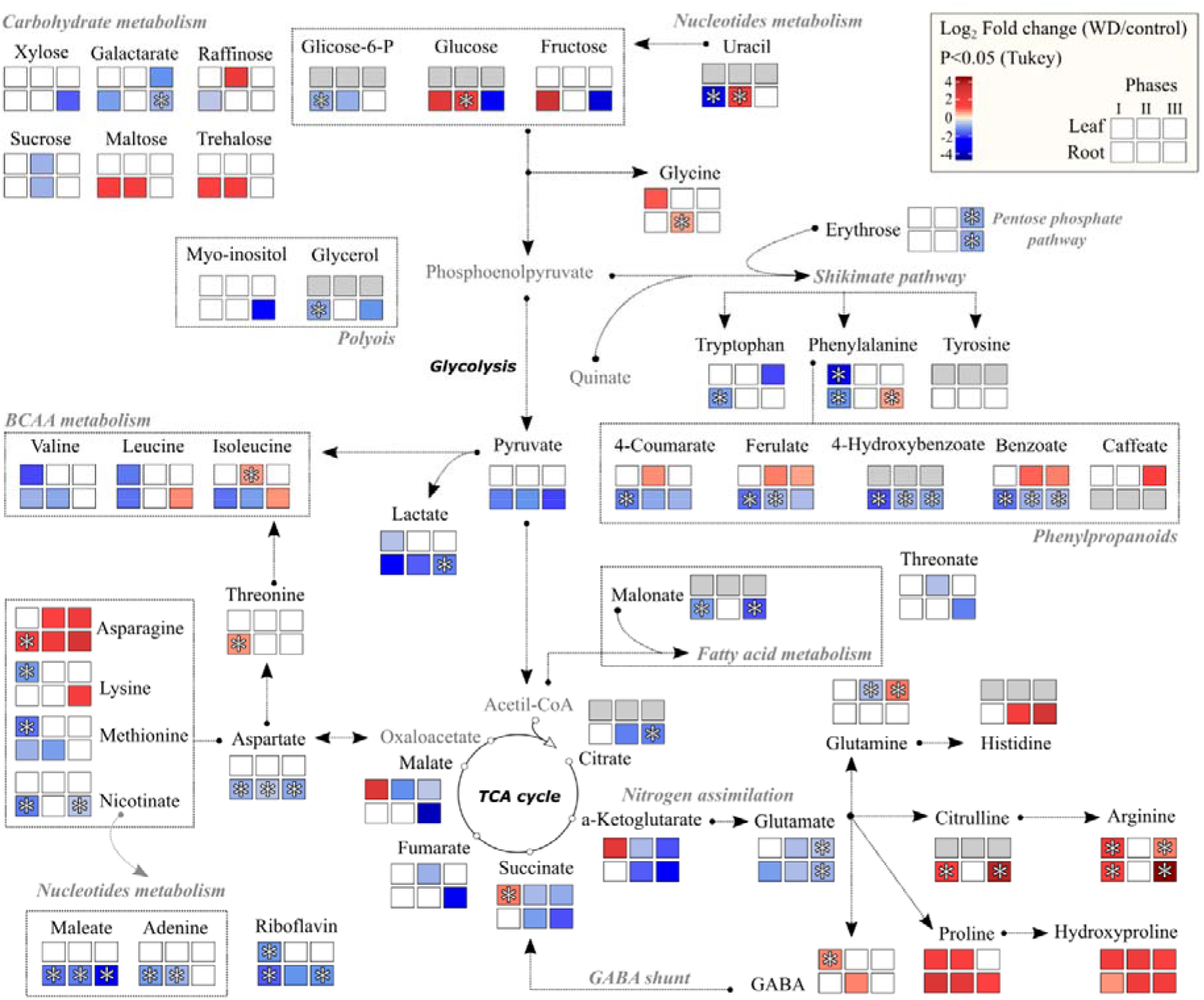
Metabolic map containing the heatmaps of relative intensities (*n* = 4) of metabolites significantly altered in leaves (Valencia sweet orange) and roots (Rangpur lime) of citrus plants under water deficit in phases I, II and III. Asterisks indicate alterations found only in plants grafted on Rangpur lime. Metabolites not found are in gray color.

Differential sugar partitioning and use were found in plants on Rangpur lime (**Fig. 6**), with low investment in osmoprotection being associated with high availability of sugars for root growth in phase I (**Fig. 3d**). As root uracil levels were reduced in phase I and increased in phase II and pyrimidine metabolism is closely linked to the carbohydrate metabolism (Mainguet et al., 2009; Kafer et al., 2004), our data suggest a strong modulation of source-sink relationship in Rangpur lime associated with enhanced root growth rate (**Fig. 3d**). In addition, root adenine levels were reduced at phases I and II (**Fig. 8**), which is a nitrogenous molecule directly associated with the energetic metabolism (Zrenner et al., 2004).

Shikimic acid pathway was differently modulated in Rangpur lime roots and leaves under water deficit in phase I. Low root tryptophan levels indicate an active metabolism of many secondary metabolites, such as auxin (IAA) and melatonin (Köhl, 2016; Tzin and Galili, 2010). While IAA acts in cell expansion, melatonin has a role on the antioxidant responses to abiotic stresses (Zhang et al., 2015; Emenecker and Strader, 2020). In drought tolerant plants, increased IAA biosynthesis was associated with an up-regulation of indole-3-acetaldehyde oxidase, a key enzyme in the tryptophan degradation pathway (Gong et al., 2010; Pustovoitova et al., 2004). In the other rootstocks, tryptophan was increased, as already reported for Sunki mandarin (Sousa et al., 2022). In addition, decreases in all phenylpropanoids (**Figs. 6 and 8**) may indicate loosening of the cell wall, which may also favor root growth (Wang et al., 2016). In fact, Rangpur lime showed increases in root growth and less sugar accumulation in phase I (**Figs. 3d and 6**), which suggests increased energy consumption to maintain root growth.

Only plants grafted on Rangpur lime showed arginine and citrulline accumulation in roots and arginine accumulation in mature leaves under water deficit (**Figs. 6 and 7**). Arginine is not only one of the main forms of organic nitrogen storage and transport in plants (Xia et al., 2014; Frémont et al., 2013) but also a precursor for nitric oxide (NO), a signaling molecule associated with increased root growth under water deficit (Silveira et al., 2016; Winter et al., 2015). This result suggests that sensitivity of Rangpur lime roots is likely associated with NO. Although arginine and citrulline have accumulated in roots and asparagine in young leaves at the recovery period for all rootstocks, their contents were particularly high in Rangpur lime (**Figs. 4 and 5**). Interestingly, leaf arginine accumulation was found only on Rangpur lime after rehydration (**Fig. 7**). Asparagine has a high N/C ratio and is one of the main forms of storage and transport of organic nitrogen in plants (Winter et al., 2015) and might contribute to plant recovery under carbon-limiting conditions (Fujiki et al., 2001). We know nitrogen assimilation is inhibited because of low levels of carbohydrate during the water deficit, ensuring energy savings while avoiding the degradation of amino acids (Köhl, 2016). Studies with *Arabidopsis thaliana* showed that arginine synthesis was upregulated after rehydration (Batista-Silva et al., 2019). If the recovery of metabolism depends on new protein synthesis (Lyon et al., 2016), the resume of plant growth after water deficit would be favored on Rangpur lime, which would justify its higher shoot growth rate during the rehydration (**Fig. 3c**).

## Conclusion

Citrus rootstocks presented differential responses and strategies under water deficit. As reported in previous studies (Santana-Vieira et al., 2016; Sousa et al., 2022), Sunki mandarin rootstock seemed to invest more energy in protective compounds and had impaired plant growth, while Swingle citrumelo resisted against drought and presented minor metabolic changes. Rangpur lime responded earlier to the initial variation of water availability, prioritizing root growth rather than shoot growth, a physiological strategy to balance water uptake and loss when such resource is limiting. In addition, Rangpur lime showed greater recovery capacity, resuming shoot growth during rehydration. Metabolic adjustments in plants grafted on Rangpur lime agreed with strategies to cope with water deficit, such as changes in nucleotide, shikimate pathway and specific amino acids (e.g. arginine, citrulline and asparagine). These changes represent differential partitioning of sugars, chemical signaling IAA and NO and increase in nitrogen use efficiency.

## Acknowledgements

The authors acknowledge the financial support provided by the National Council for Scientific and Technological Development (CNPq, Brazil; Grant #401104/2016-8 to RVR, #302460/2018-7 to RVR, and #311345/2019-0 to ECM) and the São Paulo Research Foundation (FAPESP, Brazil; Grant #2016/11906-9 to RVR), as well as the scholarships to SFS (FAPESP, Grant #2015/14817-4), MTM (FAPESP, Grant #2016/02199-7 and #2018/09834-5) and APD (FAPESP, Grant #2017/15939-1). We are also grateful to the assistance provided by Dr. Daniela F. S. P. Machado during starch analyses. Research supported by the Brazilian Biorenewables National Laboratory (LNBR/CNPEM/MCTIC) during the use of the Metabolomics open access facility (MET-21585 and MET-22973).

## Author contributions

SFS, ECM and RVR designed the experiments; SFS and MTM executed the experiments, collected data and carried out all statistical analyses; JAA performed high-throughput gas chromatography with time-of-flight mass spectrometer; CPC, APD and CC assisted in the treatment and interpretation of metabolic dataset; SFS and RVR interpreted the entire dataset; SFS and RVR wrote the first draft; and all authors edited the manuscript and approved the final version.

## Conflict of interest

The authors declare no conflict of interest.

## Supporting Information

**Table S1.** Metabolomic profiling of citrus roots (raw data).

**Table S2.** Metabolomic profiling of citrus leaves (raw data).

**Table S3.** Metabolomic profiling of citrus roots (normalized data).

**Table S4.** Metabolomic profiling of citrus leaves (normalized data).

**Table S5.** Metabolites identified in roots and leaves passing the filters.

**Table S6.** Metabolome data used in roots heatmap. Log_2_ FC (WD/control). The sample and metabolite order follow the heatmap clustering.

**Table S7.** Metabolome data used in leaves heatmap. Log_2_ FC (WD/control). The sample and metabolite order follow the heatmap clustering.

**Fig. S1.** Experimental design: (a) phases of water deficit and time-points of sampling and evaluation; (b) General aspect of plants at the onset and at the end of the experimental period, highlighting mature leaves and young leaves developed in new shoots.

**Fig. S2.** Variation of main trunk height (ΔTH, a-c), main trunk diameter (ΔTD, d-f), dry mass of previously formed leaves (LDM, g-i), dry mass of main trunk (TDM, j-l) of Valencia orange scions grafted on Swingle citrumelo (a,d,g,j), Rangpur lime (b,e,h,k) or Sunki mandarin (c,f,i,l) in each phase of water deficit. Control refers to well-watered plants while WD to water-stressed plants. Symbols represent mean values (*n* = 4) ± SD. * and ** indicate statistical differences between water treatments at 3<BF_10_<20 and BF_10_>20, respectively. The first shaded area represents phase I of water deficit while the second shaded area represents phase III (rehydration). Between them, phase II is shown.

**Fig. S3.** Dry mass of new shoots (SDM, a-c) and roots (RDM, d-f) of Valencia orange scions grafted on Swingle citrumelo (SC, a,d), Rangpur lime (RL, b,e) or Sunki mandarin (SM, c,f) in each phase of water deficit cycle. Control refers to well-watered plants while WD to water-stressed plants. Symbols represent mean values (*n* = 4) ± SD. * and ** indicate statistical differences between water treatments at 3<BF_10_<20 and BF_10_>20, respectively. The first shaded area represents phase I of water deficit while the second shaded area represents phase III (rehydration). Between them, phase II is shown.

**Fig. S4.** Starch content in young leaves (a-c) and roots (d-f) of Valencia orange scions grafted on Swingle citrumelo (SC, a,d), Rangpur lime (RL, b,e) or Sunki mandarin (SM, c,f) in each phase of water deficit cycle. Control refers to well-watered plants while WD to water-stressed plants. Boxplots consider the median, the 25th and 75th percentiles, the error bars indicate range within 1.5 IQR (*n* = 4). * and ** indicate statistical differences between water treatments at 3<BF_10_<20 and BF_10_>20, respectively.

## Notes

### Competing Interest Statement

The authors have declared no competing interest.

